# Alternative (backdoor) androgen production and masculinization in the human fetus

**DOI:** 10.1101/365122

**Authors:** Peter J O’Shaughnessy, Jean Philippe Antignac, Bruno Le Bizec, Marie-Line Morvan, Konstantin Svechnikov, Olle Söder, Iuliia Savchuk, Ana Monteiro, Ugo Soffientini, Zoe C Johnston, Michelle Bellingham, Denise Hough, Siladitya Bhattacharya, Natasha Walker, Panagiotis Filis, Paul A Fowler

## Abstract

Masculinization of the external genitalia in humans is dependent on formation of 5α-dihydrotestosterone (DHT) through both the canonical androgenic pathway and an alternative (backdoor) pathway. The fetal testes are essential for canonical androgen production but little is known about the synthesis of backdoor androgens despite their known critical role in masculinization. In this study, we have measured plasma and tissue levels of endogenous steroids in second trimester human male fetuses using multi-dimensional and high-resolution mass-spectrometry. Results show that androsterone is the principal backdoor androgen in the fetal circulation and that DHT is undetectable (<1ng/ml). Backdoor pathway intermediates are found primarily in the placenta and fetal liver with significant androsterone levels also in the fetal adrenal. Backdoor intermediates, including androsterone, are mostly undetectable in the fetal testes. This is consistent with transcript levels of enzymes involved in the backdoor pathway (SRD5A1, AKR1C2/4, CYP17A1), as measured by qPCR. These data identify androsterone as the predominant backdoor androgen in the human fetus and show that it is formed primarily in non-gonadal tissue with placental progesterone the likely substrate. Masculinization of the human fetus depends, therefore, on androgen synthesis by both the fetal testes and non-gonadal tissues leading to DHT formation at the genital tubercle. Our findings provide, for the first time, a solid basis to explain why placental insufficiency is associated with disorders of sex development in humans

## Introduction

The male external genitalia are the most common site of congenital abnormalities in the human, with up to 0.8% of male births affected (1;2). The most frequent of these abnormalities is hypospadias, which is characterized by abnormal opening of the urethra on the ventral side of the penis. Normal masculinization of the fetus is dependent upon androgen secretion by the testis and androgens act initially during a critical masculinization programming window to ensure normal male reproductive development (3). In humans, male-specific morphological differentiation and subsequent growth of the genital tubercle/penis begins around 10-12 weeks of gestation (4;5) and continues through the second and third trimesters (6;7). The etiology of hypospadias is probably multifactorial but it is likely that altered androgen exposure during the second trimester is a significant factor (8).

During masculinization testosterone acts directly to stabilize the mesonephric (Wolffian) ducts and to induce testis descent. However, it is conversion of testosterone to the more potent 5α-dihydrotestosterone (DHT) at the target organ which leads to masculinization of the external genitalia (9). In humans, testosterone is synthesized in the testicular Leydig cells through the canonical Δ^5^ pathway shown in Fig 1 (10;11). More recently, however, it has been reported that an alternative pathway to DHT formation exists which does not require testosterone as an intermediate. This alternative, “backdoor”, pathway (Fig 1) was first described in the testes of pouch young marsupials (12) and a similar pathway has since been reported in the prepubertal mouse testis (13) and the fetal human testis (14). The importance of the backdoor pathway to normal human development was initially unclear, but studies by Fluck *et al* (14) have shown that disordered sex development (DSD) will arise if the pathway is disrupted. In one individual with mutations in the aldo-keto reductase enzyme AKR1C2 and in another family with an added mutation in *AKR1C4*, there was failure of normal masculinization. Importantly, the consequences of these mutations in the backdoor pathway are similar to those seen in individuals with mutations in the canonical pathway (15). These data demonstrate that both the canonical and backdoor pathways are essential for normal fetal masculinization.

**Figure 1.**
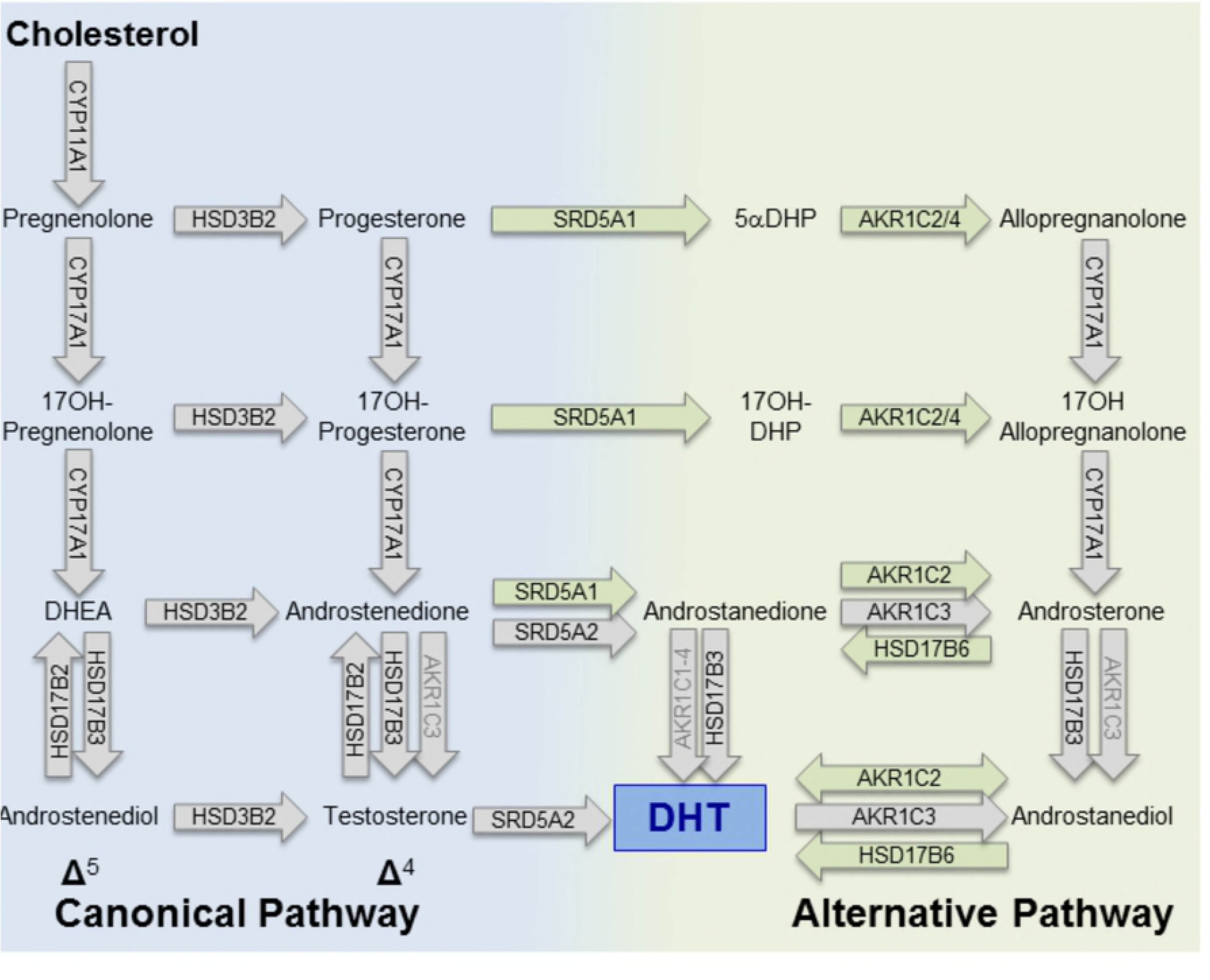
Canonical and alternative (backdoor) pathways of dihydrotestosterone synthesis. The canonical pathway has potential Δ^4^ and Δ^5^ sub-pathways. The enzymes which catalyze each step are indicated within the arrows. Enzymes written in black on a grey arrow are essential components of the canonical pathway and some appear in both canonical and backdoor pathways (e.g. CYP17A1). Enzymes written in black on a green background are specific to the backdoor pathway. Enzymes in grey text will carry out the described conversion but they may not be the principal enzyme involved. Other enzymes, not shown, may also be involved in components of the backdoor pathway (61). Abbreviations: DHEA, dehydroepiandrosterone; Androstenediol, androst-5-ene-3β,17β-diol; Androstanedione, 5α-androstane-3,17-dione; Androstanediol, 5α-androstane-3α, 17β-diol;1 5αDHP, 5α-dihydroprogesterone; 17OHDHP, 17α-hydroxydihydroprogesterone; DHT, dihydrotestosterone.

Currently, the accepted model for masculinization is that circulating DHT, formed via the backdoor pathway in the fetal testis (14;15), is important for virilization alongside circulating testosterone. At present, however, our understanding of the regulation of human fetal masculinization is seriously hindered because we do not know which steroids are present in the male fetal circulation or fetal tissues, what the concentrations of these steroids are and which tissues are involved in their metabolism. This means that the circulating levels of DHT and potential substrates for DHT synthesis at the target organ, from either the canonical or backdoor pathways, remain unknown in the human fetus.

In this study we have measured (i) concurrent levels of fetal plasma and tissue steroids by hyphenated mass spectrometric tools (ii) transcript levels of critical enzymes in the backdoor pathway in human fetal tissues and (iii) canonical and backdoor androgen synthesis by the human fetal testis *in vitro*. Our results show that high levels of intermediates in the backdoor pathway are present in the human fetal circulation and that androsterone is the major circulating backdoor androgen. Crucially, the results also show that the fetal testis contains only low levels of backdoor androgens and DHT and that androsterone is likely to be formed in non-gonadal tissues largely through metabolism of placental progesterone and adrenal dehydroepiandrosterone (DHEA).

## Results

### High levels of intermediates in the backdoor pathway are present in male fetal plasma but DHT is undetectable

Overall levels of steroids involved in the synthesis of DHT in male fetal plasma (from cardiac puncture ex-vivo) are shown in Fig 2. As expected, all steroids in the canonical Δ^4^ and Δ^5^ pathways were present in the fetal circulation. Noticeably, the data also shows that intermediates in the backdoor pathway were present at levels comparable to the Δ^5^ pathway, with the backdoor pathway clearly going from progesterone through 5α-dihydroprogesterone (5αDHP), allopregnanolone, 17α-hydroxyallopregnanolone to androsterone (Fig 2). Circulating DHT, however, was not detectable in any of the 42 fetuses (<1ng/ml). The Δ^5^ steroids pregnenolone, 17α-hydroxypregnenolone and DHEA were present at the highest levels in the fetal circulation and these steroids probably come from the fetal adrenal gland (16;17). Levels of progesterone were also high and were likely to be derived principally from the placenta (18) (and see below). The key initial intermediates in the backdoor pathway, 5αDHP and allopregnanolone, were present in the fetal circulation at similar concentrations to progesterone (means: progesterone 258ng/ml, 5αDHP 135ng/ml, allopregnanolone 243ng/ml). The principal Δ^4^ and backdoor androgens detectable in most samples were androstenedione, testosterone and androsterone, all potential substrates for DHT synthesis. Most forms of 5α-androstanediol were undetectable in fetal plasma (including 5α-androstane- 3β,17β-diol), although 5α-androstan-3α,17β-diol (labelled androstanediol in Figs 1 and 2), which is a potential substrate for DHT synthesis, was detectable in 10/42 samples (Fig 2). Etiocholanolone, a metabolite of androstenedione, was also present in most samples (Supplementary Fig 1). Levels of most steroids did not change over the course of the second trimester, with the exceptions of testosterone which declined significantly (P<0.048) and androstenediol which increased (P<0.004) over the same period (Supplementary Fig 2). Maternal smoking had no significant effect on fetal plasma steroid levels. A full list of steroids measured by GC-MS/MS in human male fetal plasma is shown in Supplementary Table 3.

**Figure 2.**
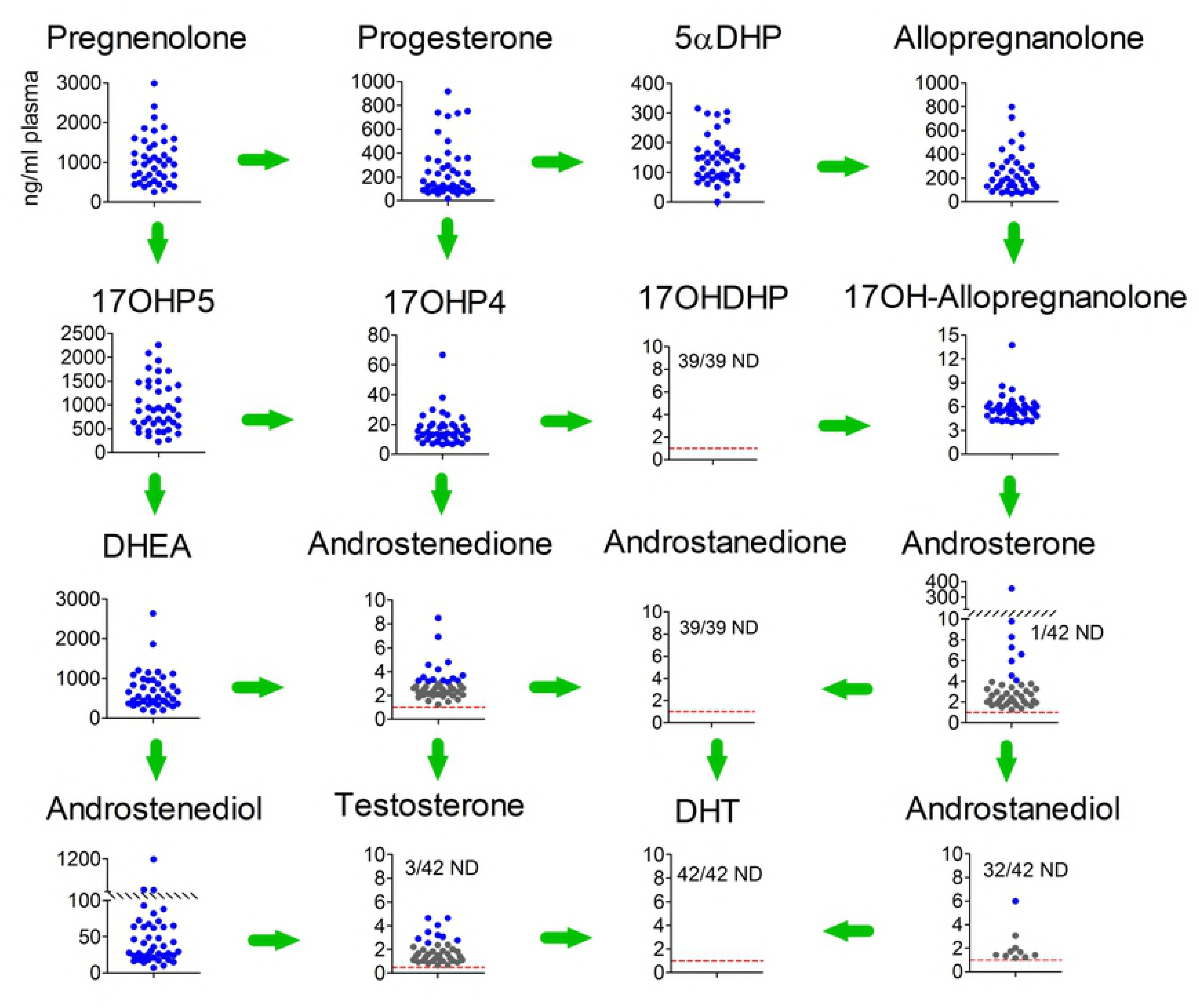
Concentrations of both canonical and backdoor steroids in the plasma of human male fetuses during the second trimester. Steroid levels from 39 or 42 individual fetuses are shown in each graph, arranged as in the pathways shown in Fig 1. The number of samples which were non-detectable (ND) for each steroid are shown and, where appropriate, the limit of detection (LOD) is shown as a red dotted line. Data shown in grey was above the LOD but below the formal LOQ which means that the quantified data shown for these samples is less reliable than for data show in blue which was above the LOQ. The plasma used in these studies was from fetuses aged between 12 and 19 weeks. Abbreviations used are the same as those in Figure 1.

### Backdoor intermediates are present primarily in the placenta and fetal liver with androsterone primarily in the fetal adrenal

Levels of major Δ^4^, Δ^5^ and backdoor androgens in placenta, fetal liver, fetal adrenal and fetal testis are shown in Fig 3. Note that 17α-hydroxylated intermediates were not measured in this study while matrix effects meant that the 5α-reduced androgen, androstanedione, was not detectable. The placenta contained high levels of progesterone with lower amounts of 5αDHP and allopregnanolone. The backdoor androgens, androsterone and androstanediol were detectable in about half the placentas while the Δ^4^ steroids androstenedione and testosterone were detectable in most placentas. DHT was also detectable in about half the placental samples (Fig 3). The major steroids detectable in the fetal liver were progesterone, allopregnanolone and DHEA. Low levels of androsterone and DHT were also detectable in most fetal livers while androstanediol was detectable in about half the samples (Fig 3). The fetal adrenals contained high levels of pregnenolone, progesterone and DHEA and androsterone was present in most adrenals. Testosterone was present in about half the adrenals but other steroids were not detectable (Fig 3). The fetal testes contained high levels of pregnenolone and testosterone with lower levels of progesterone and androstenedione. The backdoor intermediates 5αDHP and allopregnanolone were detectable in 6 and 9 testes respectively (out of 25) but androsterone was not detectable and androstanediol was only detectable in one testis. Low levels of DHT were detectable in 5 testes. To confirm the low/undetectable levels of 5α-reduced androgens in the fetal testis, testicular extracts from a further 6 fetuses were measured by GC-MS/MS (Table 2) to increase sensitivity (see Materials and Methods). These fetuses were aged 15-19 weeks and the levels of pregnenolone, DHEA, progesterone, androstenedione and testosterone were generally within the range shown in Fig 3 (all data in Fig 3 is from fetuses aged 13 or 14 weeks). In those cases measured by GC-MS/MS the backdoor steroids allopregnanolone, androstanedione, androsterone and androstanediol were detectable at very low levels in all 6 testes. Low levels of DHT were also detected in all testes (Table 2).

**Figure 3.**
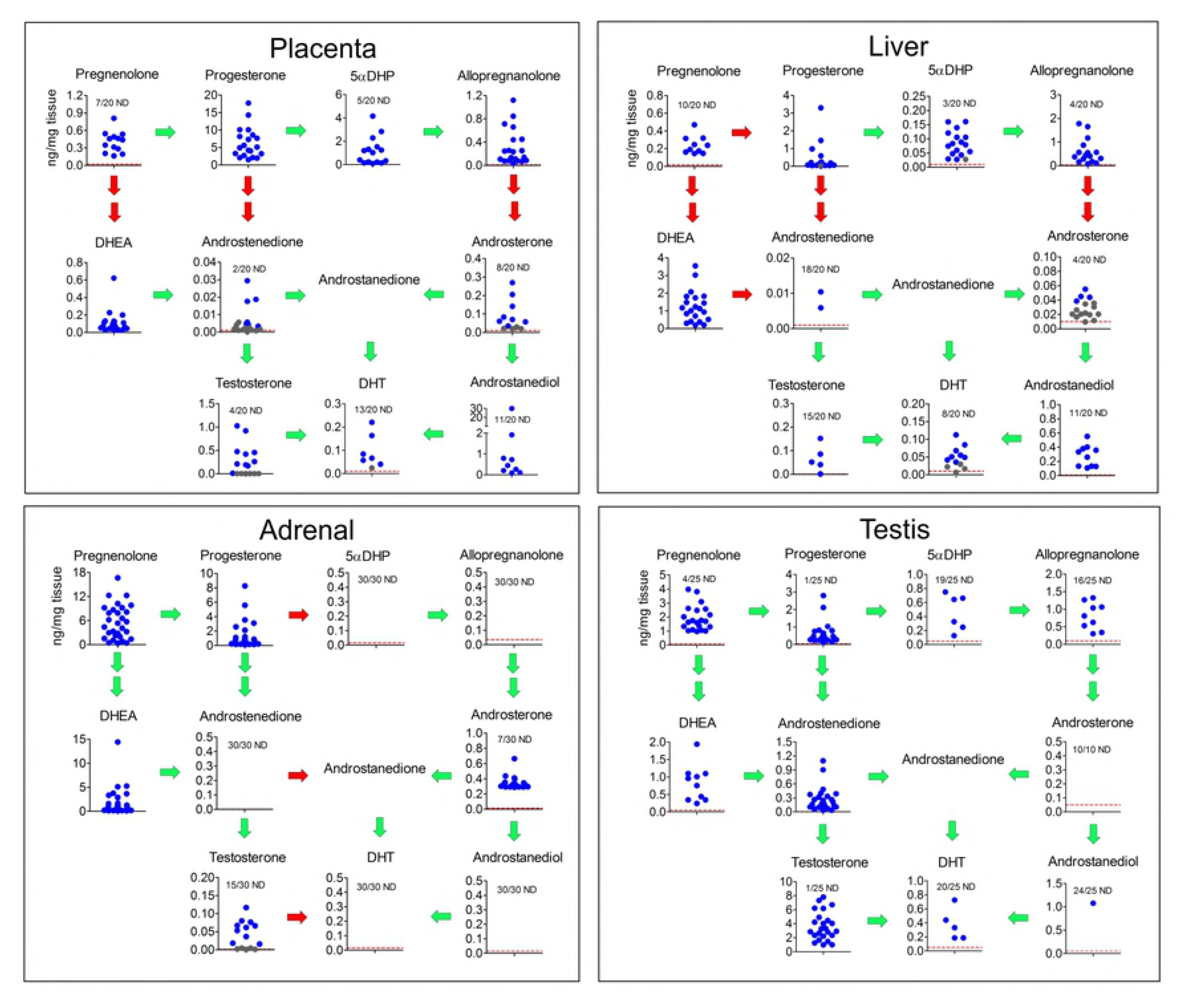
Fetal tissue levels of steroid intermediates involved in the canonical and backdoor synthesis of DHT. Tissue levels of steroids from the placenta and fetal liver (n=20; placentas and fetuses were from the same pregnancies), fetal adrenal (n=30) and fetal testis (n=10 (for DHEA and androsterone) or 25 (for all other steroids)) are shown as individual points in each graph and arranged in the pathways shown in Fig 1. Levels of 17α-hydroxylated intermediates were not measured in this part of the study. The number of samples which were non-detectable (ND) for each steroid are shown and, where appropriate, the limit of detection (LOD) is shown as a red dotted line. Data shown in grey was above the LOD but below the formal LOQ which means that the quantified data shown for these samples is less reliable. The LOD for each sample (in ng/mg tissue) depended on the mass of tissue extracted and the lines drawn are based on the average mass of each tissue used. Green arrows indicate that the relevant enzymes are detectable (as mRNA transcripts) in that tissue while red arrows indicate that the presumed enzyme is not detectable (based on data in Figure 4). Abbreviations used are the same as those in Figure 1.

To determine whether human fetal testes produce backdoor steroids under hormonal stimulation, dispersed fetal testicular cells were incubated with or without hCG for 24 h and the steroids produced were measured by GC-MS/MS. In the canonical pathway 17α-hydroxyprogesterone, DHEA and androstenedione were detectable in most samples as was pregnenolone at low levels (Supplementary Fig 3). The presence of hCG had a stimulatory effect on DHEA levels. DHT was detected in one control culture. No backdoor androgens, or intermediates in their synthesis, were detectable in any testicular cell cultures.

### Enzymes associated with the backdoor pathway are predominantly expressed in non-gonadal tissues

The critical entry-point to the backdoor pathway is through 5α-reduction of progesterone or 17α-hydroxyprogesterone by SRD5A1. The highest levels of *SRD5A1* expression in the second trimester fetus were in the liver, with significant, but lower expression in the placenta, testis and genital tubercle (Fig 4). *SRD5A2* expression was only consistently detectable in the genital tubercle. The placenta and fetal liver are considerably larger than the other organs measured in this study (Table 1) and, in terms of *total* fetal transcript levels, therefore, these tissues have about 1,000 times greater *SRD5A1* expression than the testis. AKR1C2 is specific to the backdoor pathway and is critical for human fetal masculinization (14). Mean *AKR1C2* transcript levels were highest in the fetal liver and fetal testis (Fig 4) although, taking tissue mass into account, liver and placenta each have ∼200 times more total AKR1C2 transcript than the fetal testes. There was also significant *AKR1C2* expression in the genital tubercle which is likely to be important for local DHT synthesis from androsterone (Fig 1). AKR1C4 is the other backdoor enzyme which may be required for masculinization (14) and transcripts were only consistently detected in the fetal liver (Fig 4). Expression of *HSD17B6* was highest in the testis with transcripts also consistently detected in the adrenal and placenta and lower expression in the genital tubercle (Fig 4). *HSD17B3* was expressed at similar levels in the testis and liver, with very low or undetectable expression in the placenta, adrenal and genital tubercle (Fig 4). *AKR1C3* showed the highest level of expression in the liver with low but detectable expression in other tissues. The cytochrome P450 enzyme CYP17A1 is essential in both canonical and backdoor pathways of androgen synthesis (Fig 1). Predictably, expression was high in both fetal adrenal and fetal testis, with the mean adrenal level about 6 times that of the testis (Fig 4). Expression was very low or undetectable in the genital tubercle, liver and placenta. The HSD3B2 enzyme is essential for *de novo* androgen synthesis and highest levels of *HSD3B* were in the placenta with lower expression in the testis and adrenal (Fig 4).

**Table 1.** Fetal tissue weights during the second trimester

| <u>Tissue</u> | <u>Tissue weight (mg)*</u> | <u>Fold-differences vs</u><br><u>combined testis weights</u> | <u>Reference</u> |
| --- | --- | --- | --- |
| Liver | 700 – 14,800 | 117 – 220 | our data and (59) |
| Testis (combined) | 6 – 67 | 1 – 1 | our data |
| Adrenal (combined) | 50 – 1,300 | 8 – 19 | our data |
| Genital tubercle | 3 – 98 | 0.5 – 1.5 | our data |
| Placenta | 14,000 – 115,000 | 2,300 – 1,716 | (60) |

\* Range of tissue weights from the start to the end of the second trimester in the human

**Table 2.**
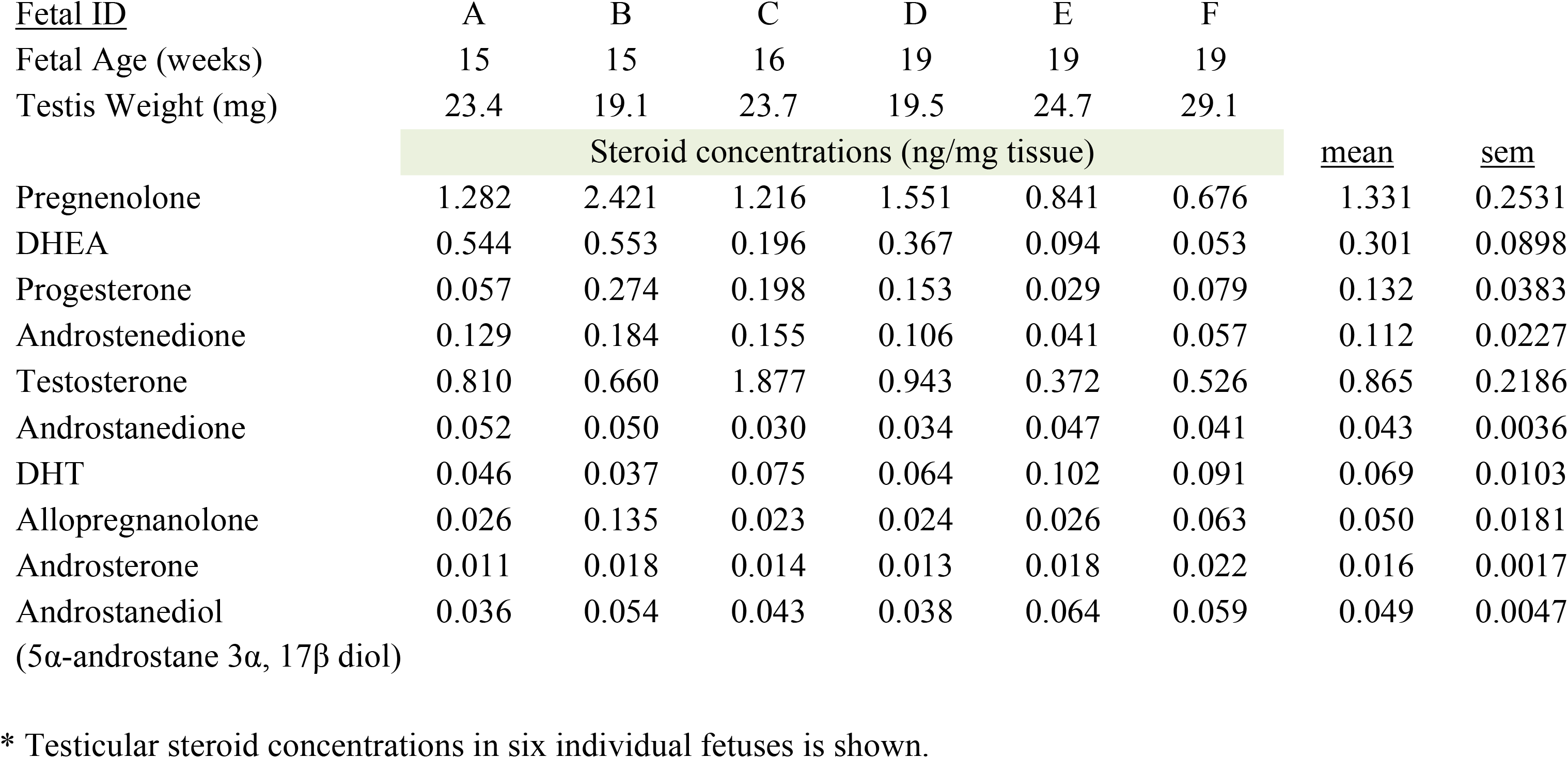
Steroids in fetal testis measured by GC-MS/MS*

**Figure 4.**
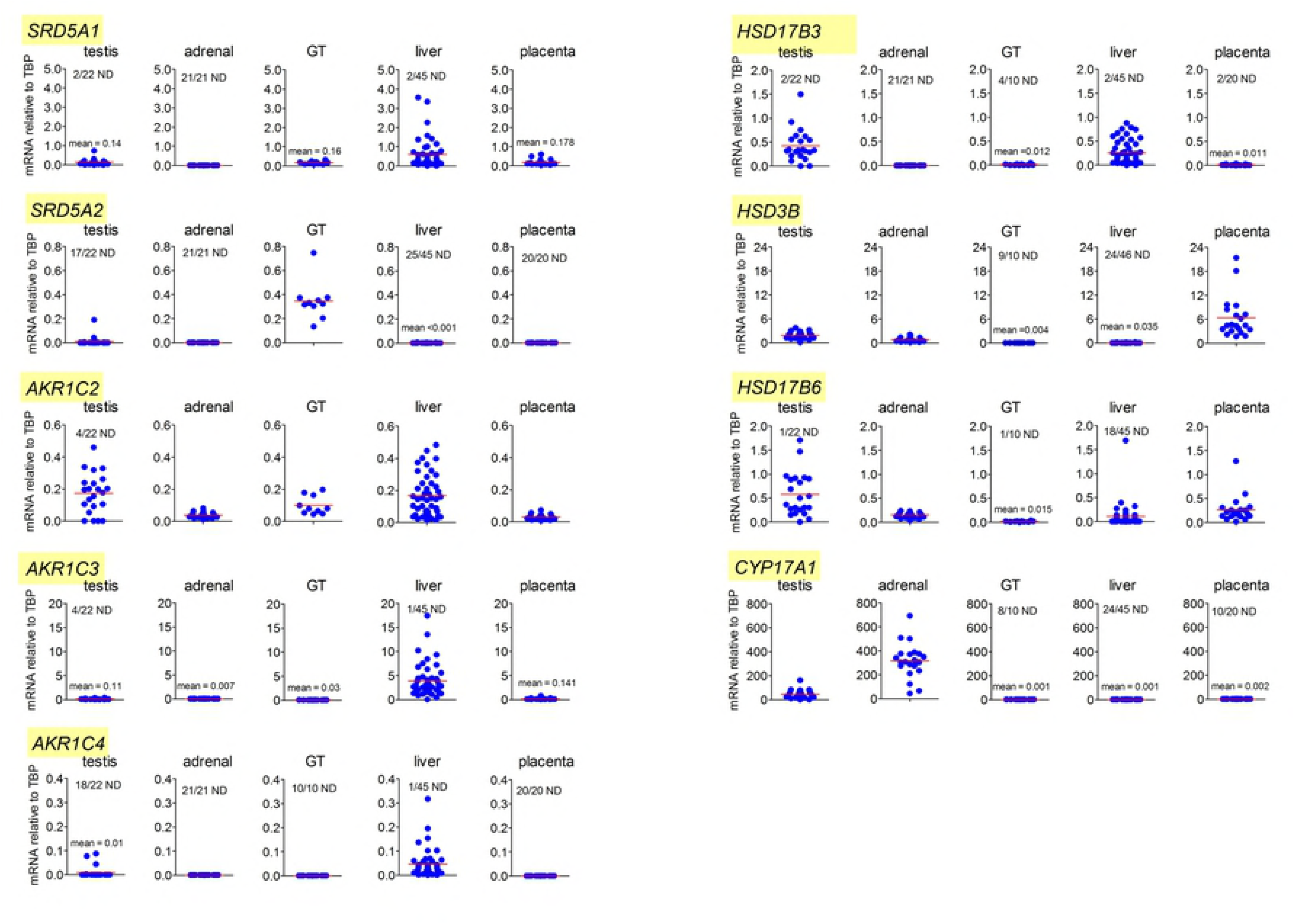
Fetal tissue levels of mRNA transcripts encoding enzymes involved in the synthesis of DHT through the backdoor pathway. Data shows levels of transcripts in testis (n=22), adrenal (n=21), liver (n=45), genital tubercle (n=10), liver (n=46) and placenta (n=20) from individual male fetuses during the second trimester. Transcript levels have been measured relative to the housekeeping gene *TBP*. For each transcript the Y axis has been maintained constant for all 5 tissues so that direct comparison of transcript levels can be made. The number of non-detectable (ND) samples is shown on each graph. The horizontal red bar indicates mean expression and, in cases where levels are very low, the mean is also provided in text on the graph.

## Discussion

Masculinization of the fetus is dependent on the action of testosterone at the Wolffian ducts and on the action of DHT at the external genitalia (19). The process of masculinization at the external genitalia starts in the early second trimester and the most intense phase of penile growth also occurs later in the second trimester (20). This is a critical period for normal masculinization, therefore, and it was assumed, until recently, that growth of the external genitalia was solely dependent on DHT formed in the target organ through 5α-reduction of testis-derived testosterone. However, the recent demonstration that the alternative, backdoor pathway to DHT synthesis is also required for normal human fetal male development (14) has shown that the process is a complex interaction between different steroidogenic pathways. In this study we now show for the first time that androsterone is the major circulating backdoor androgen in the human male fetus and that most circulating androsterone comes from non-gonadal tissues that use placental progesterone (or its metabolites) as substrate. Masculinization depends, therefore, not only on the fetal testes, but also on other, non-gonadal tissues.

The backdoor pathway of androgen synthesis depends, initially, on progesterone formed from pregnenolone and in the fetus the major *de novo* sources of pregnenolone are the adrenal and the testis. In both tissues, however, pregnenolone is metabolized predominantly through the Δ^5^ pathway to DHEA because pregnenolone is bound with a significantly higher affinity by human CYP17A1 than by HSD3B2 (21). In addition, HSD3B activity is likely to be relatively low compared to CYP17A1 in both tissues, based on transcript levels (this study and (22;23)). Both tissues contain progesterone, at levels similar to those reported previously (24), but the tissue concentrations, particularly in the testis, are not markedly greater than those in the liver which lacks significant HSD3B expression and does not have the capacity to synthesize progesterone *de novo* (25). This means that most progesterone in fetal human tissues and circulation is likely to come from the placenta which has been shown previously to secrete progesterone directly into the fetal circulation at high levels (26;27), similar to those reported here.

Metabolism of progesterone to 5αDHP depends on SRD5A1 and transcript levels of this enzyme are highest in the liver, with lower levels also present in the placenta and testis. This is consistent with earlier *in vivo* studies showing high relative 5α-reductase activity in the fetal liver (28). Tissue levels of 5αDHP are highest in the placenta, however, which probably reflects the high concentration of progesterone substrate in this tissue. Within the backdoor pathway, the conversion of 5αDHP to allopregnanolone by AKR1C2 and/or AKR1C4 has been shown to be critical for masculinization (14) and the highest consistent tissue concentrations of allopregnanolone are found in the placenta and fetal liver (Fig 3). High concentrations in the fetal liver are likely to be a reflection of high AKR1C2 and AKR1C4 and, as the fetal liver lacks *CYP17A1* expression, allopregnanolone will not be further metabolized. High expression of *AKR1C2* and *AKR1C4* in the fetal liver is likely to be another reason why 5αDHP levels are low in this tissue. The placenta does not contain high levels of *AKR1C2*, and *AKR1C4* is absent, but substrate levels for the enzymes are high and, like the liver, there will be little further metabolism of allopregnanolone via CYP17A1. In the fetal testis 5αDHP and allopregnanolone levels were undetectable in about two thirds of samples but were at a significant level in the remaining samples and might be expected to contribute to circulating steroid concentrations in these fetuses. The fetal adrenals do not express *SRD5A1/2* and would not be expected to contribute to fetal 5αDHP or allopregnanolone production. Taking tissue mass into account, the liver and placenta are likely to be the major sites of 5αDHP and allopregnanolone production in the second trimester fetus. Placental allopregnanolone production would also be consistent with the increasing plasma levels in pregnant women during gestation (29). Finally, it should be noted that 50% or more of any steroids secreted into the fetal circulation by the placenta will go directly to the fetal liver through the umbilical vein and so most fetal 5α-reduction or 3α-hydroxylation of placental progesterone is likely to take place in the fetal liver.

Once formed, there is a high affinity between allopregnanolone and human CYP17A1 (30) but the only tissues which express the enzyme are the fetal testes and adrenals. The high circulating levels of allopregnanolone in the fetus, and the relatively low 17α-hydroxyallopregnanolone and androsterone levels would suggest that conversion by CYP17A1 is limiting in the fetus. Tissue androsterone levels were highest in the adrenals which is consistent with high *CYP17A1* transcript levels and, since the adrenals lack *SRD5A1* and appear unable to make 5α-reduced steroids, the substrate for this reaction must come from circulating allopregnanolone. It should be noted that this androsterone was mostly the sulfated form which is a reflection of the high SULT2A1 levels in the human fetal adrenal (17). The presence of androsterone in the fetal adrenals at the end of the first trimester has also been reported previously (22). In contrast to the adrenals, androsterone levels in the fetal testis were either undetectable or extremely low. Differences between the fetal testis and adrenal are likely to be a reflection of the lower *CYP17A1* in the testis, much greater vascularization in the fetal adrenal for allopregnanolone substrate provision (31) and, possibly, greater competition from other substrates in the testis. Interestingly, the placenta also contained detectable androsterone in most samples. There is some debate about whether the human placenta expresses significant CYP17A1 and associated 17α-hydroxylase and C_17-20_ lyase activity (32;33) and our data shows that transcripts are either absent or are present only at very low levels. This suggests that most placental androsterone must be derived from adrenal DHEA although the possibility exists for some *de novo* synthesis. Given the relative sizes of the fetal organs involved and the tissue concentrations, the placenta and the fetal adrenals are likely to be, by far, the major sources of backdoor androsterone production in the fetus.

When the backdoor pathway was shown to be essential for human fetal masculinization, it was suggested that DHT is synthesized via this pathway in the fetal testes and released into the circulation (14;15). This now appears very unlikely because DHT is undetectable, or present at very low levels, in testes from most fetuses and because circulating DHT levels are undetectable (<1ng/ml). These results are consistent with earlier studies which either failed to detect DHT in the human fetal testis (34;35) or found very low levels (36). DHT can be metabolized to androstanediol by all AKR1 enzymes, and AKR1C2 in particular (11;37), and given the high levels of AKR1 enzyme transcripts in the fetal liver, it is likely that any DHT released into the fetal circulation is rapidly metabolized. Our results show, therefore, that the major circulating backdoor androgen in the fetus is androsterone which is present at similar levels to testosterone. It remains to be shown whether the genital tubercle can convert androsterone to DHT but the tissue expresses high levels of *AKR1C2*, as well as detectable *AKR1C3*, and AKR1C2 can catalyze both oxidation and reduction steps required for androsterone conversion to DHT (Figs 1 and 5). [A schematic diagram of the proposed pathways involved in backdoor androgen synthesis in the human fetal male is shown in Fig 5.

**Figure 5.**
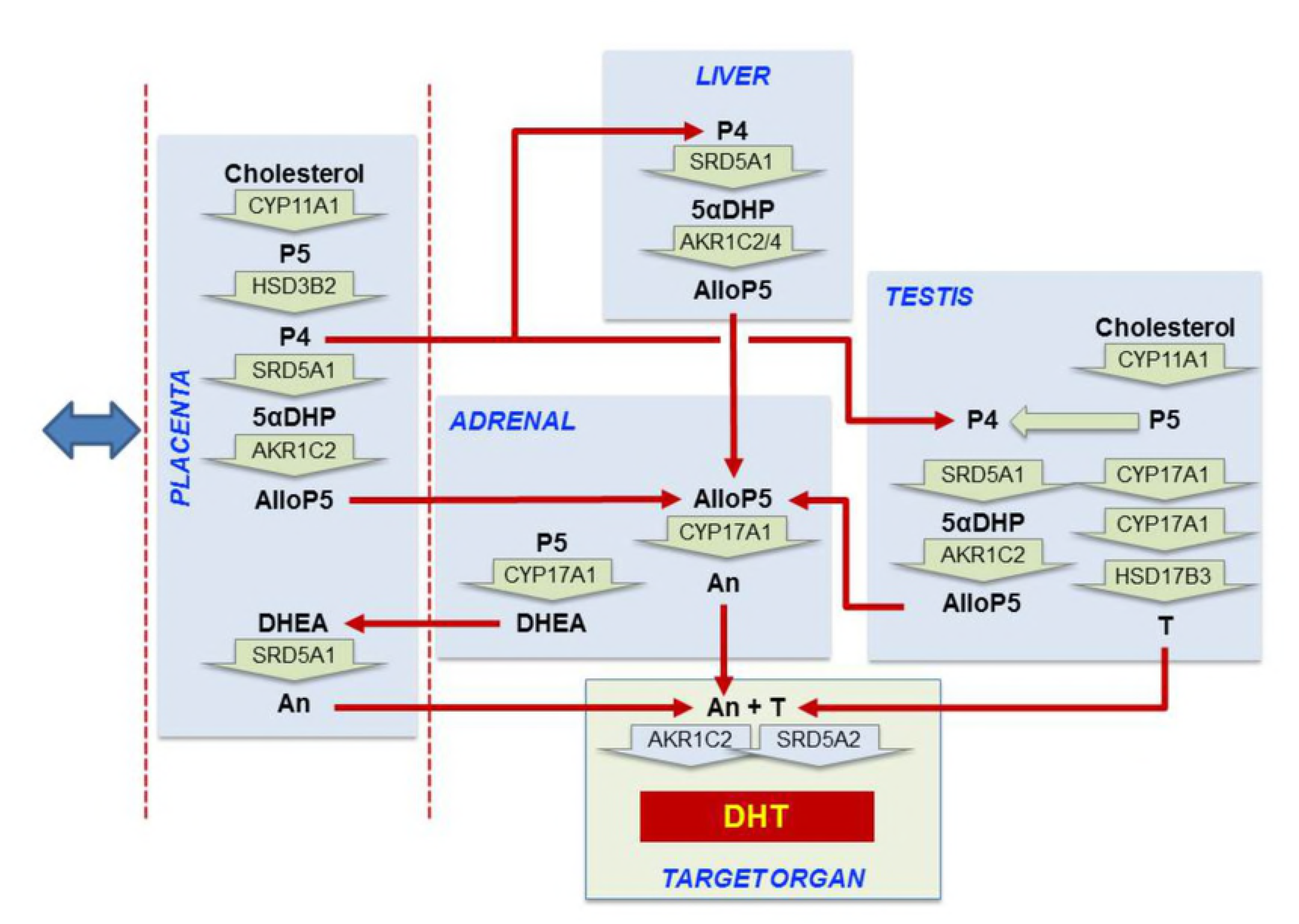
Proposed steroidogenic pathways leading to androsterone (An) synthesis and masculinization in the second trimester human male fetus. Steroid hormone conversion is shown by wide green arrows with the converting enzymes written within the arrow. Red arrows show transport between organs in the fetal circulation. The blue double-headed arrow indicates that exchange is also taking place between the placenta and the maternal circulation. Most circulating progesterone in the fetal circulation is likely to come from the placenta and this will be reduced to 5α-dihydroprogesterone (5αDHP) by SRD5A1 in the placenta, fetal liver and fetal testis with the fetal liver likely to be the major site. Allopregnanolone (AlloP5) production by AKR1C2 is also most likely to occur in the placenta and fetal liver because the substrate is present in those tissues and they express the highest total levels of enzyme transcript. Some conversion may also occur in the testis. Significant levels of androsterone are only detectable in the placenta and adrenal so they are the likely source of the circulating steroid. The adrenal lacks other intermediates in the backdoor pathway so AlloP5 must come from other tissues. The placenta lacks CYP17A1 so androsterone production is likely to depend on adrenal DHEA as substrate. Testosterone (T) from the fetal testes also acts as an essential substrate for DHT synthesis at the external genitalia. Abbreviations: P5, pregnenolone; P4, progesterone; 5αDHP, 5α-dihydroprogesterone;1 AlloP5, allopregnanolone; DHEA, dehydroepiandrosterone; An, androsterone; T, testosterone; DHT, 5α-dihydrotestosterone.

In the DSD cases described by Flück et al (14) the 46,XY patients carried hypomorphic mutations in *AKR1C2* and *AKR1C4* or *AKR1C2* alone. From data reported here, reduced fetal AKR1C2 activity would be expected to affect backdoor pathways in the liver and testis reducing production of allopregnanolone. In addition, loss of enzyme activity in the genital tubercle may also affect production of DHT at the target organ. The potential involvement of placental AKR1C2 in the reported 46,XY DSD patients is also of interest. The placenta develops from both maternal and fetal cells and it is not known whether placental AKR1C2 is of fetal or maternal origin. It is of note, however, that two 46,XY individuals who are known heterozygotes for a mutation in *AKR1C2* show divergent phenotypes with one showing a normal male phenotype and the other DSD (14). The mother of the affected individual is also heterozygous for a mutation in *AKR1C2* and if placental AKR1C2 is of maternal origin then this would be expected to affect backdoor steroid production in the placenta. Unfortunately the genotype of the mother of the unaffected heterozygous individual is not available.

Further strong evidence for a non-testicular pathway of backdoor androgen synthesis comes from patients with P450 oxidoreductase (POR) or 21-hydroxylase deficiency who show 46,XX virilization. It is likely that virilization in these individuals *in utero* is due to excessive backdoor androgen synthesis which is occurring in the absence of testes (38-40). It has been postulated that this increase in fetal backdoor androgen production comes from the fetal adrenals (41;42). The lack of *SRD5A1* and intermediates in the backdoor pathway in normal fetal adrenals make this unlikely, however, unless the condition itself increases adrenal SRD5A1 activity (e.g. through increased adrenal stimulation by ACTH). Our data would suggest that the increased fetal adrenal 17α-hydroxyprogesterone seen in these conditions acts initially as substrate for backdoor androgen production through 5α-reduction in other tissues, probably the fetal liver.

If the placenta is a critical component of the fetal backdoor androgen pathway, as suggested by this data, then it has implications for our understanding of the regulation of masculinization and DSD. It is now established that placental insufficiency, associated with intrauterine growth restriction (IUGR), is associated with abnormalities in development of the male external genitalia, and hypospadias in particular (43;44). Severe forms of placenta-mediated IUGR start during the first trimester (43) and could interfere, therefore, with all aspects of fetal masculinization. It has been suggested that DSD associated with placental dysfunction could be due to reduced placental hCG production (44) but other studies have shown that maternal hCG levels tend to be increased in placental insufficiency (43). In contrast, maternal progesterone levels are reported to be reduced during IUGR (45), suggesting placental steroidogenesis is affected. If the placenta is central to fetal backdoor androgen production, as we suggest, then altered placental steroidogenesis may lead directly to abnormalities in masculinization.

Studies in the 1950s/60s by Jost and others showed that androgen is required for masculinization of the external genitalia (reviewed in (46)). Later studies showed that testicular testosterone must be converted to DHT at the target organ to induce masculinization (47). Most recently, it has been shown that the normal process of masculinization in the human depends on two separate pathways of androgen synthesis, the canonical and backdoor pathways (14). We now report that the backdoor pathway in the human fetus is driven by placental progesterone production which acts as substrate for androsterone synthesis, primarily in non-gonadal tissues. This means that our current understanding of the endocrine control of masculinization in the human fetus is that it is mediated through circulating testosterone and androsterone and is dependent on a complex interaction and exchange between the testes and non-gonadal tissues, particularly the placenta.

## Methods

### Sample collection

Three sources of human fetal tissues were used in this study: 1) In Aberdeen, human fetuses between 11 and 21 weeks of gestation and classified as normal at scan were collected from women over 16 years of age undergoing elective termination (17) Information about maternal smoking during pregnancy was available for most fetuses. Fetuses were transported to the laboratory within 30 minutes of delivery, weighed, crown-rump length recorded, and sexed. Blood samples from a total of 42 male fetuses were collected by cardiac puncture *ex vivo* and plasma was stored at −80°C. Tissues were snap-frozen in liquid N_2_ and then stored at −80°C. 2) In Stockholm, human fetal testes were obtained for *in vitro* incubation studies after elective termination of pregnancy at 10–12 weeks of gestation. 3) Additional fetal livers and placentas were provided by the MRC/Wellcome-Trust funded Human Developmental Biology Resource (HBDR, http://www.hdbr.org). Available fetal and maternal characteristics relevant to samples used in different parts of this study are shown in Supplementary Table 1.

Collections of fetal material in Aberdeen, Stockholm and by HBDR were respectively approved by the NHS Grampian Research Ethics Committees (REC 04/S0802/21), the Regional Ethics Committee of Stockholm (EPN dnr 2014/1022-32) and the relevant Research Ethics Committees, in accordance with the relevant UK Human Tissue Authority (HTA; www.hta.gov.uk) Codes of Practice. Written, informed, maternal consent was received from participants prior to inclusion in the study.

### RNA Extraction, Reverse Transcription and Real-Time PCR

Total RNA was extracted from frozen fetal tissues either using TRIzol (Life Technologies, Paisley, UK) (48) or using Qiagen AllPrep kits (Qiagen, Manchester, UK). Reverse transcription, primer design and real-time PCR were carried out as previously described (49;50) and the primers used are shown in Supplementary Table 2. RNA which is free of genomic DNA contamination is required to amplify *SRD5A1* because of the presence of a processed pseudogene in the genome and this was carried out using RNAeasy Plus Micro-columns (Qiagen Ltd, Manchester, UK) followed by DNase treatment (DNA-free, Life Technologies, Paisley, UK). To normalize data, Normfinder was used to identify the most stable housekeeping genes in each tissue using the housekeeping genes and primers described earlier (51). The best combinations of housekeeping genes varied between tissues and so, to allow comparison of transcript expression between tissues, *TBP* (TATA box binding protein) was used as housekeeping gene for all samples since it was the most consistently stable transcript across all tissues (51;52). Some steroid enzyme transcript data (*SRD5A1*, *SRD5A2*, *CYP17A1* and *HSD3B* in liver (25) and *HSD17B3* and *CYP17A1* in testis (53)) has been reported previously relative to different housekeeping genes or external standards. This data is shown again here relative to *TBP* to allow comparisons between tissues and additional fetal liver samples have been included in the reported data. Fetal sex was confirmed by PCR of *ZFX* and *SRY* using genomic DNA and primers described in Supplementary Table 2 (54).

### Isolation and incubation of human fetal testicular cells

Isolated fetal testes were treated with collagenase type I (Sigma Chemicals Co., St. Louis, USA) (1 mg/ml for 35 min at 37°C) and then disrupted mechanically. The testicular cells were collected by centrifugation at 300 g for 7 min, washed in Hank’s balanced salt solution and resuspended in DMEM-F12 supplemented with 1 mg/ml BSA, 100 IU/ml penicillin and 100 μg/ml streptomycin. For incubation, 100 μl of a suspension containing 1.5×10^5^ cells/ml was plated onto 96-well plates (Falcon, Franklin Lakes, NJ, USA) in the presence or absence of hCG (10 ng/ml) to stimulate Leydig cell activity. Cells were incubated for 24 hours at 37° C under 5% CO_2_ and steroid levels in the culture media were measured by GC-MS/MS as described below.

### Steroid extraction and quantification

Methods used to extract and profile steroid levels by GC-MS/MS in fetal plasma and culture media have been described elsewhere in detail (22;55;56). Briefly, fetal plasma samples (50 µl) or culture media samples (200 µl) were spiked with internal standards (stable isotope-labeled steroid analogues) and an enzymatic deconjugation of phase II metabolites was performed (Sulfatase, Sigma S9626 (100 U/ml) and β-Glucuronidase, Sigma G8132, (5 KU/ml)). Steroids were extracted twice with diethyl ether and a ChromP SPE cartridge was used for initial purification. Androgens and estrogens were separated by liquid/liquid partitioning with n-pentane and were further purified on a silica SPE cartridge. The androgen fraction was derivatized with MSTFA/TMIS/DTE and 2 µl of each extract was injected onto a Scion 436 gas chromatograph coupled to a Scion TQ triple quadrupole mass spectrometer (Bruker, Fremont, CA, USA). Electron ionization (70 eV) was used and two diagnostic signals (SRM acquisition mode) were monitored for identification and quantification of the targeted compounds. Levels of circulating testosterone shown in detail here have been reported previously as mean levels (57).

Extraction and quantification of tissue steroid levels by LC-HRMS have been described previously (17). Tissue samples (25-65mg) from placenta, liver and adrenals or whole testes (8-20mg, one per fetus) were initially processed to isolate RNA/DNA/protein using Qiagen Allprep kits as above and steroids were then extracted from the column-eluants following addition of internal standards (stable isotope-labeled steroid analogues) (17). Following extraction, steroids were separated by reverse-phase liquid chromatography on an Accucore™ PFP column with trimethyl silane (TMS) endcapping (50 mm, 2.1 mm, 2.6 μm, Thermo Fisher Scientific) and using an UltiMate 3000 RSLCnano autosampler/pump (Thermo Fisher Scientific, Waltham, MA, USA). Steroid were then ionized by electrospray in positive mode and signals were recorded on a Q Exactive Orbitrap (Thermo Fisher Scientific) mass spectrometer. Sulfated steroids were detected in a second analysis after negative electrospray ionization. Levels of sulfated and non-sulfated DHEA have been combined in the reported results as have sulfated and non-sulfated androsterone. The fetal testes used in this part of the study were extracted in two batches and recovery of steroid sulfates in the first batch was poor so testicular DHEA and androsterone levels have only been reported for the second batch of 10 samples. For the second batch of testicular tissue an additional extraction step using chloroform/n-butanol was used to improve steroid sulfate extraction efficiency. As reported for other LC-MS methods (58), the electrospray ionization efficiency can be low for 5α-reduced androgens which results in relatively high LOQ and LOD values for these compounds. In a separate study, therefore, testicular steroids from an additional 6 fetuses were extracted and separation and quantification of steroids in these samples was carried out by GC-MS/MS as described above for plasma samples.

### Statistics

Data were checked for normality and normalized by log-transformation as appropriate. Plasma steroid data was analyzed by 2-factor ANOVA with fetal age and maternal smoking as the factors. Correlations between steroid data and age were analyzed using Pearson’s correlation coefficient. The effect of hCG on steroid secretion by the fetal testes was analyzed by t-test.

## Author contributions

PJOS and PAF designed the study; M-LM, AM, US, ZCJ, NW, IS, DH and PF carried out the research; PJOS, PAF, KS, DH, OS, JPA, BLB, MB and SB analyzed data; POS and PAF wrote the paper.

## Acknowledgements

We thank Ms Linda Robertson, Ms Margaret Fraser, Ms Samantha Flannigan and the staff at Grampian NHS Pregnancy Counselling for their expert assistance and help. The study was supported by the following grants: Chief Scientist Office (Scottish Executive, CZG/4/742 (PAF & PJOS); NHS Grampian Endowments 08/02 (PAF, SB & PJOS) and 15/1/010 (PAF, PF, US, PJOS); the Glasgow Children’s Hospital Research Charity Research Fund, YRSS/PHD/2016/05 (NW, MB, PJOS & PAF); the European Community’s Seventh Framework Programme (FP7/2007-2013) under grant agreement no 212885 (PAF) Medical Research Council Grants MR/L010011/1 (PAF & PJOS) and MR/K501335/1 (MB, PAF & PJOS) and the Kronprinsessan Lovisas Foundation, “Stiftelsen Gunvor och Josef Anérs”, the “Stiftelsen Jane och Dan Olssons” and the “Stiftelsen Tornspiran” (KS & OS).

## References

1. Ahmed SF, Dobbie R, Finlayson AR, Gilbert J, Youngson G, Chalmers J, et al. Prevalence of hypospadias and other genital anomalies among singleton births, 1988-1997, in Scotland. Arch Dis Child Fetal Neonatal Ed. 2004;89: F149–F151.

2. Nelson CP, Park JM, Wan J, Bloom DA, Dunn RL, Wei JT. The increasing incidence of congenital penile anomalies in the United States. J Urol. 2005;174: 1573–1576.

3. Welsh M, Saunders PT, Fisken M, Scott HM, Hutchison GR, Smith LB, et al. Identification in rats of a programming window for reproductive tract masculinization, disruption of which leads to hypospadias and cryptorchidism. J Clin Invest. 2008;118: 1479–1490.

4. Overland M, Li Y, Cao M, Shen J, Yue X, Botta S, et al. Canalization of the Vestibular Plate in the Absence of Urethral Fusion Characterizes Development of the Human Clitoris: The Single Zipper Hypothesis. J Urol. 2016;195: 1275–1283.

5. Li Y, Sinclair A, Cao M, Shen J, Choudhry S, Botta S, et al. Canalization of the urethral plate precedes fusion of the urethral folds during male penile urethral development: the double zipper hypothesis. J Urol. 2015;193: 1353–1359.

6. Welsh M, Macleod DJ, Walker M, Smith LB, Sharpe RM. Critical androgen-sensitive periods of rat penis and clitoris development. Int J Androl. 2010;33: e144–e152.

7. Macleod DJ, Sharpe RM, Welsh M, Fisken M, Scott HM, Hutchison GR, et al. Androgen action in the masculinization programming window and development of male reproductive organs. Int J Androl 2010;33: 279–287.

8. Thankamony A, Pasterski V, Ong KK, Acerini CL, Hughes IA. Anogenital distance as a marker of androgen exposure in humans. Andrology 2016;4: 616–625.

9. Wilson JD, George FW, Griffin JE. The hormonal-control of sexual development. Science 1981; 211: 1278–1284.

10. Fluck CE, Miller WL, Auchus RJ. The 17, 20-lyase activity of cytochrome p450c17 from human fetal testis favors the delta5 steroidogenic pathway. J Clin Endocrinol Metab. 2003;88: 3762–3766.

11. Rizner TL, Penning TM. Role of aldo-keto reductase family 1 (AKR1) enzymes in human steroid metabolism. Steroids 2014;79: 49–63.

12. Wilson JD, Auchus RJ, Leihy MW, Guryev OL, Estabrook RW, Osborn SM, et al. 5alpha-androstane-3alpha,17beta-diol is formed in tammar wallaby pouch young testes by a pathway involving 5alpha-pregnane-3alpha,17alpha-diol-20-one as a key intermediate. Endocrinology 2003;144: 575–580.

13. Mahendroo M, Wilson JD, Richardson JA, Auchus RJ. Steroid 5alpha-reductase 1 promotes 5alpha-androstane-3alpha,17beta-diol synthesis in immature mouse testes by two pathways. Mol Cell Endocrinol. 2004;222: 113–120.

14. Fluck CE, Meyer-Boni M, Pandey AV, Kempna P, Miller WL, Schoenle EJ et al. Why boys will be boys: two pathways of fetal testicular androgen biosynthesis are needed for male sexual differentiation. Am J Hum Genet. 2011;89: 201–218.

15. Biason-Lauber A, Miller WL, Pandey AV, Fluck CE. Of marsupials and men: "Backdoor" dihydrotestosterone synthesis in male sexual differentiation. Mol Cell Endocrinol. 2013; 371: 124–132.

16. Ishimoto H, Jaffe RB. Development and function of the human fetal adrenal cortex: a key component in the feto-placental unit. Endocr Rev. 2011;32: 317–355.

17. Johnston ZC, Bellingham M, Filis P, Soffientini U, Hough D, Bhattacharya S, et al. The human fetal adrenal produces cortisol but no detectable aldosterone throughout the second trimester. BMC Med. 2018;16: 23.

18. Tuckey RC. Progesterone synthesis by the human placenta. Placenta 2005;26: 273–281.

19. Wilson JD, Griffin JE, Russell DW. Steroid 5 alpha-reductase 2 deficiency. Endocr Rev 1993;14: 577–593.

20. Gallo CB, Costa WS, Furriel A, Bastos AL, Sampaio FJ. Development of the Penis during the Human Fetal Period (13 to 36 Weeks after Conception). J Urol. 2013;190: 1876–1883.

21. Nguyen PT, Lee RS, Conley AJ, Sneyd J, Soboleva TK. Variation in 3beta-hydroxysteroid dehydrogenase activity and in pregnenolone supply rate can paradoxically alter androstenedione synthesis. J Steroid Biochem Mol Biol. 2012;128: 12–20.

22. Savchuk I, Morvan ML, Antignac JP, Gemzell-Danielsson K, Le BB, Soder O, et al. Androgenic potential of human fetal adrenals at the end of the first trimester. Endocr Connect. 2017; 6: 348–359.

23. Goto M, Piper Hanley K., Marcos J, Wood PJ, Wright S, Postle AD, et al. In humans, early cortisol biosynthesis provides a mechanism to safeguard female sexual development. J Clin Invest. 2006;116: 953–960.

24. Tapanainen J, Kellokumpulehtinen P, Pelliniemi L, Huhtaniemi I. Age-related-changes in endogenous steroids of human-fetal testis during early and mid-pregnancy. J Clin Endocrin Metab. 1981;52:98–102.

25. O’Shaughnessy PJ, Monteiro A, Bhattacharya S, Fraser MJ, Fowler PA. Steroidogenic enzyme expression in the human fetal liver and potential role in the endocrinology of pregnancy. Mol Hum Reprod. 2013;19:177–187.

26. Partsch CJ, Sippell WG, MacKenzie IZ, Aynsley-Green A. The steroid hormonal milieu of the undisturbed human fetus and mother at 16-20 weeks gestation. J Clin Endocrinol Metab. 1991;73:969–974.

27. Pasqualini JR. Enzymes involved in the formation and transformation of steroid hormones in the fetal and placental compartments. J Steroid Biochem Mol Biol. 2005;97: 401–415.

28. Stern MD, Ling W, Coutts JR, Macnaughton MC, Solomon S. Metabolism of testosterone in previable human fetuses. J Clin Endocrinol Metab. 1975;40: 1057–1065.

29. Luisi S, Petraglia F, Benedetto C, Nappi RE, Bernardi F, Fadalti M, et al. Serum allopregnanolone levels in pregnant women: changes during pregnancy, at delivery, and in hypertensive patients. J Clin Endocrinol Metab. 2000;85: 2429–2433.

30. Gupta MK, Guryev OL, Auchus RJ. 5alpha-reduced C21 steroids are substrates for human cytochrome P450c17. Arch Biochem Biophys. 2003;418:151–160.

31. Hervonen A, Suoranta H. Vascular supply of human fetal adrenals and the functional significance of the microvascular patterns. Z Anat Entwicklungsgesch 1972;136: 311–318.

32. Pion R, Jaffe R, Eriksson G, Wiqvist N, Diczfalusy E. Studies on the metabolism of C-21 steroids in the human foeto-placental unit. I. formation of a beta-unsaturated 3-ketones in midterm placentas perfused in situ with pregnenolone and 17-alpha-hydroxypregnenolone. Acta Endocrinol (Copenh). 1965;48: 234–248.

33. Escobar JC, Patel SS, Beshay VE, Suzuki T, Carr BR. The human placenta expresses CYP17 and generates androgens de novo. J Clin Endocrinol Metab. 2011;96: 1385–1392.

34. Siiteri PK, Wilson JD. Testosterone formation and metabolism during male sexual differentiation in the human embryo. J Clin Endocrinol Metab. 1974;38: 113–125.

35. Huhtaniemi IT, Korenbrot CC, Jaffe RB. hCG binding and stimulation of testosterone biosynthesis in the human fetal testis. J Clin Endocrinol Metab. 1977;44: 963–967.

36. George FW, Carr BR, Noble JF, Wilson JD. 5-Alpha-reduced androgens in the human-fetal testis. J Clin Endocrinol Metab. 1987;64: 628–630.

37. Steckelbroeck S, Jin Y, Gopishetty S, Oyesanmi B, Penning TM. Human cytosolic 3alpha-hydroxysteroid dehydrogenases of the aldo-keto reductase superfamily display significant 3beta-hydroxysteroid dehydrogenase activity: implications for steroid hormone metabolism and action. J Biol Chem. 2004;279: 10784–10795.

38. Reisch N, Idkowiak J, Hughes BA, Ivison HE, Abdul-Rahman OA, Hendon LG, et al. Prenatal diagnosis of congenital adrenal hyperplasia caused by P450 oxidoreductase deficiency. J Clin Endocrinol Metab. 2013;98: E528–E536.

39. Kamrath C, Hochberg Z, Hartmann MF, Remer T, Wudy SA. Increased activation of the alternative "backdoor" pathway in patients with 21-hydroxylase deficiency: evidence from urinary steroid hormone analysis. J Clin Endocrinol Metab. 2012;97: E367–E375.

40. Arlt W, Walker EA, Draper N, Ivison HE, Ride JP, Hammer F, et al. Congenital adrenal hyperplasia caused by mutant P450 oxidoreductase and human androgen synthesis: analytical study. Lancet 2004;363: 2128–2135.

41. Fukami M, Homma K, Hasegawa T, Ogata T. Backdoor pathway for dihydrotestosterone biosynthesis: implications for normal and abnormal human sex development. Dev Dyn. 2013;242: 320–329.

42. Auchus RJ, Miller WL. Congenital adrenal hyperplasia--more dogma bites the dust. J Clin Endocrinol Metab. 2012;97: 772–775.

43. Yinon Y, Kingdom JC, Proctor LK, Kelly EN, Salle JL, Wherrett D, et al. Hypospadias in males with intrauterine growth restriction due to placental insufficiency: the placental role in the embryogenesis of male external genitalia. Am J Med Genet A. 2010;152A: 75–83.

44. Fredell L, Kockum I, Hansson E, Holmner S, Lundquist L, Lackgren G, et al. Heredity of hypospadias and the significance of low birth weight. J Urol. 2002;167: 1423–1427.

45. Pecks U, Rath W, Kleine-Eggebrecht N, Maass N, Voigt F, Goecke TW, et al. Maternal serum lipid, estradiol, and progesterone levels in pregnancy, and the impact of placental and hepatic pathologies. Geburtshilfe Frauenheilkd 2016;76: 799–808.

46. Jost A. Hormonal factors in the sex differentiation of the mammalian foetus. Philos Trans R Soc Lond Biol 1970;259: 119–130.

47. Wilson JD. The role of 5alpha-reduction in steroid hormone physiology. Reprod Fertil Dev 2001;13: 673–678.

48. Fowler PA, Cassie S, Rhind SM, Brewer MJ, Collinson JM, Lea RG, et al. Maternal smoking during pregnancy specifically reduces human fetal desert hedgehog gene expression during testis development. J Clin Endocrinol Metab. 2008 93:619–626.

49. O’Shaughnessy PJ, Willerton L, Baker PJ. Changes in Leydig cell gene expression during development in the mouse. Biol Reprod. 2002;66: 966–975.

50. O’Shaughnessy PJ, Baker PJ, Monteiro A, Cassie S, Bhattacharya S, Fowler PA. Developmental changes in human fetal testicular cell numbers and messenger ribonucleic acid levels during the second trimester. J Clin Endocrinol Metab. 2007;92: 4792–4801.

51. O’Shaughnessy PJ, Monteiro A, Fowler PA. Identification of stable endogenous reference genes for real-time PCR in the human fetal gonad using an external standard technique. Mol Hum Reprod. 2011;17: 620–625.

52. O’Shaughnessy PJ, Monteiro A, Bhattacharya S, Fowler PA. Maternal smoking and fetal sex significantly affect metabolic enzyme expression in the human fetal liver. J Clin Endocrinol Metab. 2011;96: 2851–2860.

53. O’Shaughnessy PJ, Baker PJ, Monteiro A, Cassie S, Bhattacharya S, Fowler PA. Developmental changes in human fetal testicular cell numbers and messenger ribonucleic acid levels during the second trimester. J Clin Endocrinol Metab. 2007;92: 4792–4801.

54. Fain S, LeMay P Gender indentification of human and mammalian wildlife species from PCR amplified sex linked genes. Proc Am Acad Forensic Sci. 1995;1: 34.

55. Courant F, Aksglaede L, Antignac JP, Monteau F, Sorensen K, Andersson AM, et al. Assessment of circulating sex steroid levels in prepubertal and pubertal boys and girls by a novel ultrasensitive gas chromatography-tandem mass spectrometry method. J Clin Endocrinol Metab. 2010;95: 82–92.

56. Courant F, Antignac JP, Maume D, Monteau F, Andersson AM, Skakkebaek N, et al. Exposure assessment of prepubertal children to steroid endocrine disrupters 1. Analytical strategy for estrogens measurement in plasma at ultra-trace level. Anal Chim Acta 2007;586: 105–114.

57. Fowler PA, Filis P, Bhattacharya S, Le Bizec B, Antignac JP, Morvan ML, et al. Human anogenital distance: an update on fetal smoke-exposure and integration of the perinatal literature on sex differences. Hum Reprod 2016;31: 463–472.

58. McDonald JG, Matthew S, Auchus RJ. Steroid profiling by gas chromatography-mass spectrometry and high performance liquid chromatography-mass spectrometry for adrenal diseases. Horm Cancer 2011;2: 324–332.

59. Archie JG, Collins JS, Lebel RR. Quantitative standards for fetal and neonatal autopsy. Am J Clin Pathol. 2006;126:256–265.

60. Molteni RA. Placental growth and fetal/placental weight (F/P) ratios throughout gestation - their relationship to patterns of fetal growth. Semin Perinatol. 1984;8:94–100.

61. Penning TM, Chen M, Jin Y. Promiscuity and diversity in 3-ketosteroid reductases. J Steroid Biochem Mol Biol. 2015;151:93–101.

